# Imaging Architecture of Granulomas Induced by *Mycobacterium tuberculosis* Infections with Single-Molecule FISH

**DOI:** 10.1101/2023.02.02.526702

**Authors:** Ranjeet Kumar, Afsal Kolloli, Selvakumar Subbian, Deepak Kaushal, Lanbo Shi, Sanjay Tyagi

## Abstract

Granulomas are an important hallmark of *Mycobacterium tuberculosis* (*Mtb*) infection. They are organized and dynamic structures created by an assembly of immune cells around the sites of infection in the lungs to locally restrict the bacterial growth and the host’s inflammatory responses. The cellular architecture of granulomas is traditionally studied by immunofluorescence labeling of phenotypic surface markers. However, very few antibodies are available for model animals used in tuberculosis research, such as non-human primates and rabbits; secreted immunological markers such as cytokines cannot be imaged *in situ* using antibodies; and traditional phenotypic surface markers do not provide sufficient resolution for the detection of many subtypes and differentiation states of immune cells. Using single-molecule fluorescent *in situ* hybridization (smFISH) and its derivatives, amplified smFISH (ampFISH) and iterative smFISH, we developed a platform for imaging mRNAs encoding immune markers in rabbit and macaque tuberculosis granulomas. Multiplexed imaging for several mRNA and protein markers was followed by quantitative measurement of expression of these markers in single cells *in situ*. A quantitative analysis of combinatorial expressions of these markers allowed us to classify the cells into several subtypes and chart their distributions within granulomas. For one mRNA target, HIF-1α, we were able to image its mRNA and protein in the same cells, demonstrating the specificity of probes. This method paves the way for defining granular differentiation states and cell subtypes from transcriptomic data, identifying key mRNA markers for these cell subtypes, and then locating the cells in the spatial context of granulomas.

## Introduction

Tuberculosis has historically been and still is a major cause of death from infectious diseases in many parts of the world. Long-recognized hallmarks of tuberculosis pathology are granulomas in the lung which form around the sites of initial infection by *Mycobacterium tuberculosis* (*Mtb*). They form as tissue-resident macrophages phagocytize the pathogen while simultaneously recruiting other immune cells including dendritic cells, monocytes, neutrophils, NK cells, and T and B cells to the site of infection via the release of chemokines and cytokines (1–3). Granulomas minimize damage to larger lung tissue by locally restricting both bacterial proliferation and inflammatory immune responses (1–3).

Although signatures of the immune response to *Mtb* are present in the peripheral blood as memory T cells (4, 5) and as transcriptomic markers (6), the most intimate and long-term contact between *Mtb* and the host immune system occurs within the confines of the granuloma (3). It is therefore more informative to study interactions between the pathogen and the immune system through studies of the functional architecture of granulomas.

Since human granulomas are not readily accessible, several model animals, such as non-human primates, rabbits, mice, and zebrafish, are often used in these studies (1, 2, 7). These animal models simulate human disease to different extents and have different experimental benefits and limitations (1, 2, 7). *Mtb* infection in the non-human primate, macaque, closely mimics certain features of human disease progression, including latency and a granuloma architecture characterized by a necrotic core (1, 2, 7). Rabbits also form necrotizing granulomas similar to those found in human tuberculosis (7); whereas, *Mtb* infection in common mice strains, such as C57BL/6 or BALB/c, generally does not develop necrotic lesions, though strain C3HeB/J is reported to produce necrotic granulomas (8, 9).

Granuloma architecture has traditionally been studied using hematoxylin and eosin staining and by immunofluorescence (IF) (10–12), although recently, other approaches like multiplexed ion beam imaging by time of flight (MIBI-TOF) and *in situ* sequencing have been reported (9, 13). However, very few antibodies are available for macaques and rabbits. An additional limitation of IF is that secreted markers such as cytokines cannot be imaged *in situ* except in *ex vivo* settings where brefeldin A is used to prevent cytokine secretion. An attractive alternative is to image cells through fluorescence *in situ* hybridization (FISH) against expressed mRNAs. In a FISH variant called single-molecule FISH (smFISH), multiple small oligonucleotide probes are used against the same mRNA which allows for highly specific and sensitive detection of mRNA molecules (14, 15) (Figure 1A). In smFISH, each molecule of target RNA becomes visible as a fine fluorescent spot when viewed under high magnification objectives.

**Figure 1.**
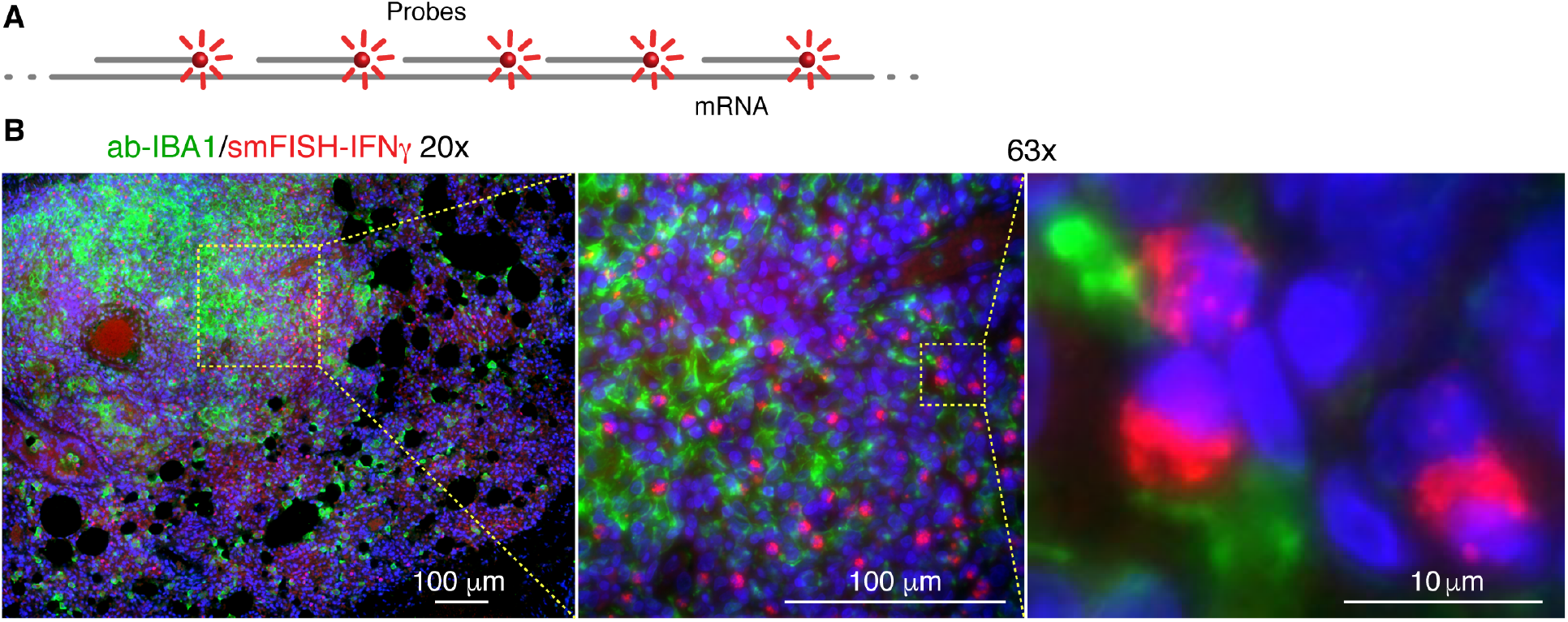
Multiplex detection of macrophages (using an antibody against IBA1) and cells that produce IFNγ (using smFISH probes against mRNA encoding IFNγ) in a lung section of a solid granuloma from a rabbit infected for four weeks by *Mtb* strain HN878.

Here we demonstrate that the functional architecture of tuberculosis granulomas can be studied in high resolution using smFISH and its variants. We imaged mRNAs encoding a number of immunological markers in rabbit and macaque tuberculosis granulomas in archived lung tissues. By combining smFISH with IF we could image both mRNA and protein markers in the same sections. For mRNA markers that are expressed at very low levels, such as surface receptors, we performed ampFISH in which pairs of interacting hairpin probes were tiled across the length of the mRNA which created amplified signals via hybridization chain reaction. Finally, we were able to image as many as eight different mRNA targets in the same tissue section, by performing iterative rounds of smFISH in which multiple targets are interrogated in successive cycles of hybridization, imaging and signal removal. Using the quantitative measures of expression of different markers in the individual cells we identified various cell types and charted their distributions in this complex tissue. These studies pave the way for transcriptome-based analysis of granuloma architecture without the limitations of IF in the model animals frequently used in tuberculosis research.

## Materials and Methods

### Lung Specimen

New Zealand female white rabbits (*Oryctolagus cuniculus*) were infected with *Mycobacterium tuberculosis* HN878 through a ‘snout-only aerosol-exposure system’, as described previously (16). This model has been demonstrated to produce progressive active TB with caseating necrotic and cavitary granulomas (16). At 4 to 8-weeks post-infection, rabbits were euthanized, and lungs were collected. Portions of the lungs were placed in 10% buffered formalin and kept in the refrigerator for at least one week. The formalin-fixed lung sections were paraffin-embedded and stored. The archived paraffin-embedded blocks were cut into 5 μm sections and used for FISH and immunofluorescence studies.

### Probes

For smFISH we designed 48 oligodeoxynucleotides probes for each mRNA using Stellaris RNA FISH probe designer tool available at (https://www.biosearchtech.com/support/tools/design-software/stellaris-probe-designer). All probe sequences and transcript IDs are provided in the Supplementary Table 1. These oligonucleotides were obtained with 3’-amino groups, pooled in equimolar concentatrtions, coupled to aminoreactive fluorophores and then purifiled by high pressure liquid chromatography as described before (17). Primary probes for iterative smFISH were obtained as a pool that contained equimolar amounts of each of the 358 oligonucelotides from IDT DNA (Supplementary Table 1). Readout probes were obtained with 3’ amino moieties and coupled to either Texas Red or Cy5 and purifed by HPLC like smFISH probes. ampFISH acceptor probes for CD3ε were prepared by click chemistry as described in Marras *et. al*. (18).

### Immunofluorescence, hybridization, and imaging

To deparaphenize and hydrate the formaldehyde fixed and parafin embeded (FFPE) tissue sections on slides, they were serially incubated for 10 min at room temperature in Xylene (twice), 100% ethanol, 90% ethanol, 70% ethanol and finally, either in hybridization wash buffer (10% formamide in 2X saline sodium citrate (SSC) buffer), or in antigen reterieval buffer (10 mM sodium citrate solution with 0.05% Tween 20, pH 6.0). For antigen retrieval, the sections were incubated in the antigen reterival buffer in a decloaking chamber, 90 °C for 40 min, followd by slow return to room temperature. In experiments where both protein and mRNAs were detected, IF was performed first followed by a secondary formaldehyde fixation (4% formaldehyde in 1X phosphate buffred saline) for 5 min.

Immunofluorescence based imaging was performed by first blocking the section with RNAse free bovine serum albumin (BSA, 5mg/ml in PBS) for 30 min, followed by incubation with IBA1 specific primary antibody, ab5076 (Abcam, Waltham, MA, USA), at 1:250 dilution. The slides were washed with PBS and incubated with an alexa 488-labelled secondary antibody, ab150129 (Abcam, Waltham, MA, USA), at 1:1000 dilution for 1h. HIF1α protein was detected using an anti HIF1α antibody directly conjugated to Dylight650 (MA5-16008,ThermoFisher Scientific, Waltham, MA, USA) for 2 hr. All IF steps were carried out at room temperature.

For smFISH, the sections were equilibrated with hybridization wash buffer and then incubated overnight in a 37 °C water bath with sufficent hybridization solution to cover the section. The hybridization solution contained 10% dextran sulfate (Sigma, St.Louis, Missouri, United States), 1 mg/ml *Escherichia coli* tRNA (Sigma), 2 mM ribonucleoside vanadyl complex (New England Biolabs, Ipswich, MA), 0.02% RNase free bovine serum albumin (Thermo Fisher Scientific, Waltham, MA)), 10% formamide, and 500 ng/ml of each probe set. After hybridization, the slides were washed twice with hybridization wash buffer at room temperature, and then mounted with 0.17 mm thick coverslips in oxygen-depleted mounting medium (0.4% w/v glucose, 2X SSC, 37 μg/ml glucose oxidase, 1% v/v of catalase suspension (both from Sigma) and 1 μg/ml DAPI).

Images were acquired using an Axiovert 200M microscope (Zeiss, Oberkochen, Germany) under the control of Metamorph software (Molecular Devices, San Jose, CA) and a Prime sCMOS camera (Teledyne Photometrics, Tucson, AZ). We recognized the granulomas by their characteristic morphology in which compressed and aggregated nuclei are surrounded by healthy tissue by visual examination in the DAPI channel and then imaged them and the surrounding tissue using a 20x objective (0.75 NA) using the scan function in the Metamorph software. Higher magnification z-stack images were acquired using a 63x objective (1.3 NA). The images were generally acquired in DIC, DAPI, TMR, Texas red and Cy 5 channels. TMR channel served as a control channel (no probes in this channel) for the detection of autofluorescence.

### Iterative smFISH

The iterative hybridization and imaging were performed on an open dish with a coverslip bottom. We placed the FFPE tissue section on a 25 mm coverslip, carried out the relatively harsher initial steps of deparaphenization, hydration, and antigen retrieval on the free-standing coverslips and then incorporated the coverslips into an open-top imaging chamber (Warner Instruments, Holliston MA). A pool of 358 primary probes (1.4 nM of each) was hybridized overnight at 37 °C in 300 μl hybridization buffer. Excess probes were removed by washing twice with hybridization wash buffer and the coverslip was mounted in the imaging chamber, which was placed on the microscope stage. Thereafter, four cycles of the following steps were performed where in each step, 300 μl of a solution was added and each step preceded with the removal of the previous solution. After each cycle the probe pair in step 2 was changed. 1. Equilibrate with readout probe hybridization solution (10% w/v ethylene carbonate, 10% dextran sulfate, 4X SSC) for 2 min. 2. Hybridize with a pair of readout probes one labeled with Texas Red and the other with Cy5 (12 nM each) in readout probe hybridization solution (sequences provided in Supplementary Table 1) for 15 min. 3. Wash (twice) with hybridization wash buffer supplemented with 0.1% Tween 20. 4. Equilibrate with glucose buffer (0.4% w/v glucose, 2XSSC), 2 min. 5. Equilibrate with oxygen-depleted mounting medium and image.

### Image Processing

For cell segmentation we used images from the DAPI channel (Figure 2), maximum intensity merges of HIF-1α smFISH and IF images (Figure 3), and maximum intensity merges of IFNγ smFISH and CD3ε ampFISH images (Figure 4). Before cell segmentation the scanned images were stitched togther in Metamorph image acquisition software (Figure 2) and z-sections were merged in ImageJ (Figure 3). These segmentation layers were incorporated into a multilayer image file (3 layers for Figure 2 and 4 layers for Figures 3 and 4) which was imported into a QuPath project. After specifying the entire image as the annotation area we used script nucleusDetectionfluo which calls for the StarDist algorythm to find nuclear boudaries (actual nuclei in Figure 2 but cells in Figures 3 and 4). The parameters were adjusted to achieve accurate segmentation of almost all objects. After detection of all cells we classified them using average pixel intensity within the area of cells for each of the individual measurement channels. From these single measurement classifiers we created composite classifiers which identified cells expressing combinations of markers. The composite images and vector graphic overlays were exported into Adobe Illustrator where final figures were created.

**Figure 2.**
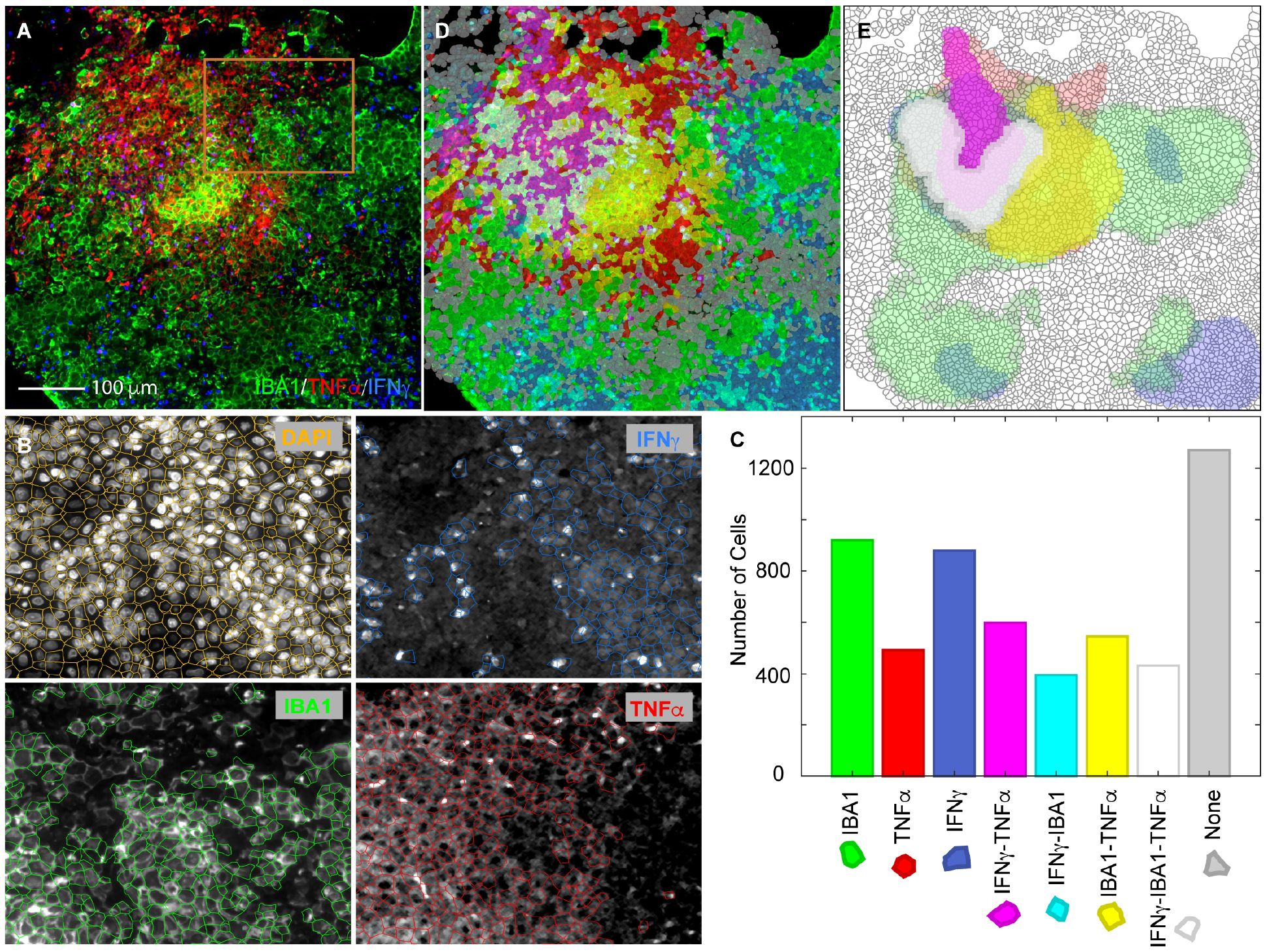
Identification of cell types based on combinatorial expression of a macrophage specific surface protein marker IBA1 and two mRNAs encoding cytokines TNFα and IFNγ in a solid *Mtb* granuloma in a rabbit lung. **A**. A color combined primary image showing IF against IBA1 (green) and smFISH against mRNAs encoding TNFα (red) and IFNγ (blue). An isotope control antibody and irrelevant smFISH probes did not yield these signals (not shown). **B**. A demonstration of the accuracy of cell detection in the section using a portion of the image in **A**indicated by an orange square. The boundaries of detected cells were overlayed on the primary images from the indicated channels in the panels shown. The cell boundaries were created using DAPI stains of nuclei and cells expressing each marker (mean pixel intensity) above a certain threshold were identified using cell intensity histograms in QuPath/StarDist. **C**. Number of cells in the image **A**that were expressing each of the three markers or their combinations. **D**. A translucent digital map of the section overlayed on the primary image in which cells were labelled with distinct colors that identify the combinations of markers expressed (color key presented in C). **E**. A density map of cells expressing various markers and their combinations. The contour maps show regions which contain more than 40 cells positive for indicated markers in a unit area of 10,000 μm^2^.

**Fig. 3.**
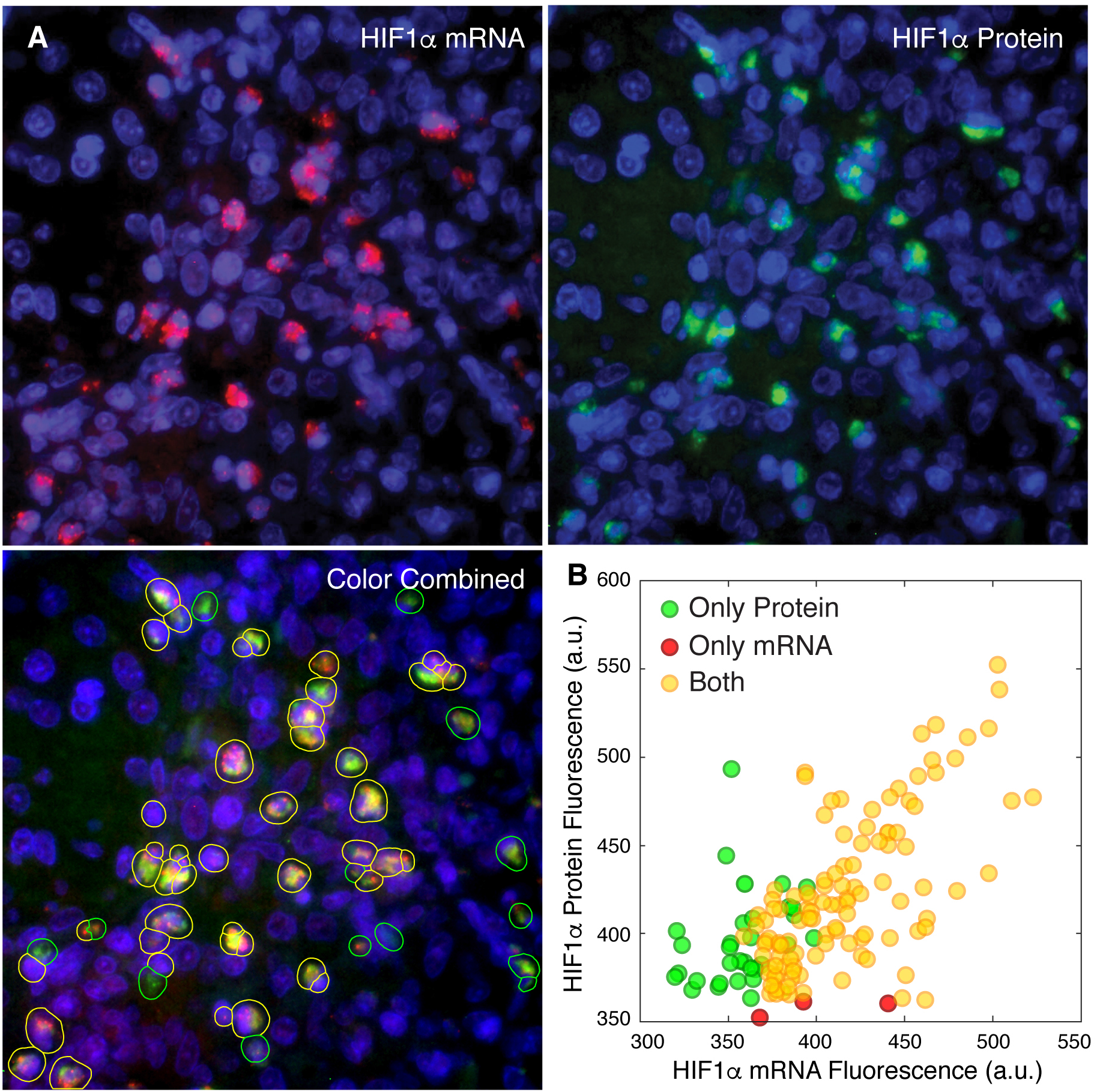
Correlated expression of a mRNA and the protein encoded by it. **A**. Three panels showing smFISH based imaging of HIF-1α mRNA (red) and antibody-based imaging of HIF-1α protein (green). Lower left panel shows the detection and classification of cells by colored overlays. **B**. Average pixel fluorescence in two channels within the boundaries of single cells from three fields in the same rabbit granuloma.

**Fig. 4.**
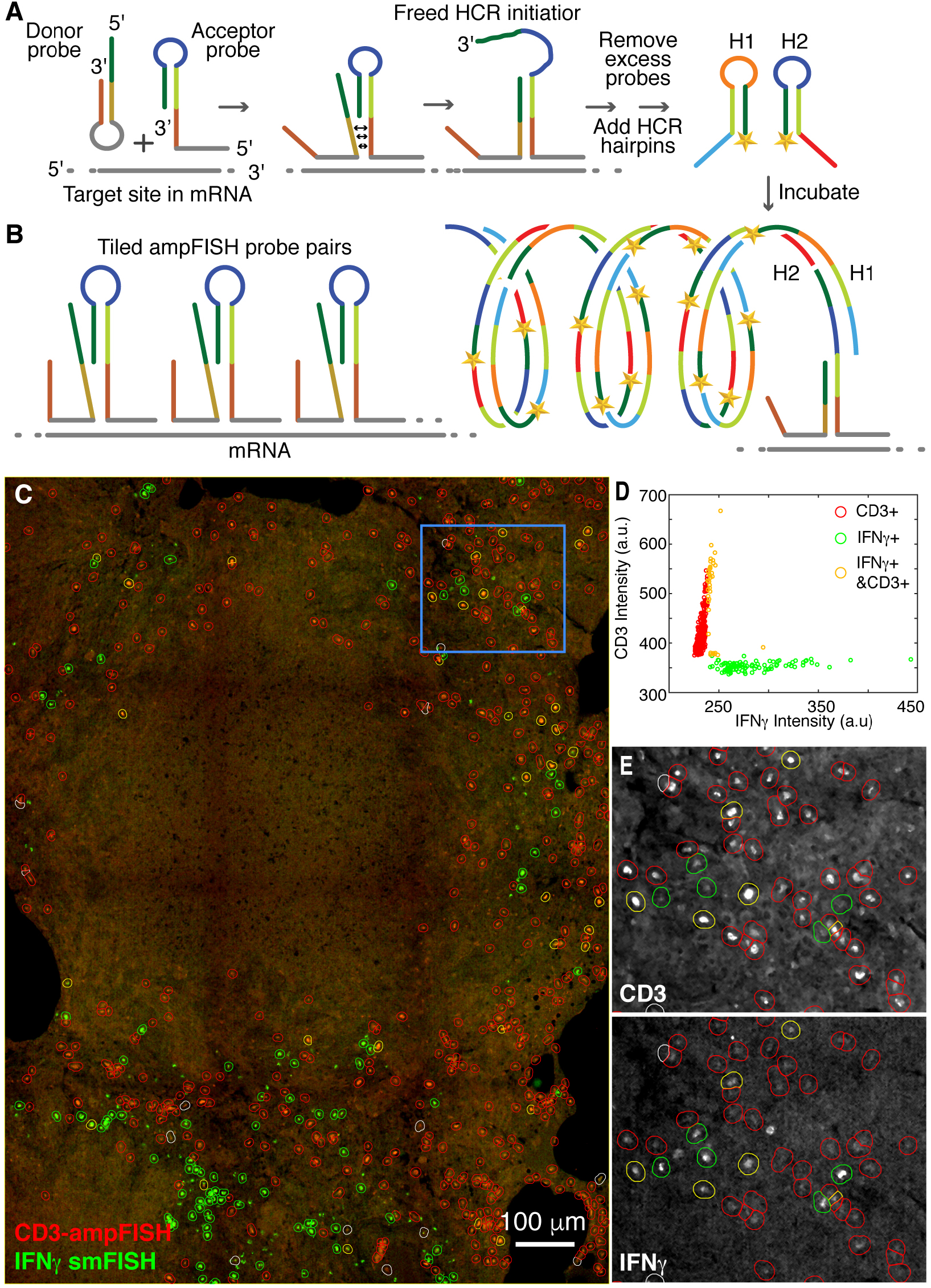
Imaging CD3ε mRNA using ampFISH and IFNγ mRNA using smFISH in within a necrotic granuloma from *Mtb* infected macaque lung. **A** and **B**. Schematic representations of ampFISH and multiple pairs of ampFISH probes tiled over the mRNA. **C** A color combined stitched image depicting Texas Red channel for IFNγ specific smFISH probes (green) and Cy5 channel for CD3 ε specific ampFISH signals (red). Cells expressing each marker (fine dotes) are identified by circles of different colors (key in D). **D**. Distribution of mean pixel intensities within areas of detected cells in each channel and the classification of three cell types based on expression of CD3ε and IFNγ individually or together. **E**. Enlargements from the area demarcated by the blue rectangle in **C**. In this pair of images, the two component channels are presented separately in grey scale to illustrate the accuracy of classification. Red circles encompass cells expressing in CD3ε channel, green circles in IFNγ channel and yellow circles in both.
.

## Results

### Challenges for RNA imaging in TB granuloma tissues

There are two major challenges in the analysis of granulomas by smFISH. First, smFISH is usually performed with high magnification objectives (63x or 100x) that yield diffraction limited spots for single mRNA molecules. However, to obtain a good overview of the distributions of different cell types in the granuloma, the imaging needs to be performed at lower magnifications which can’t resolve single-molecule spots and has lower sensitivity. To address this issue, we imaged larger regions by scanning granuloma sections with a 20x objective with a high numerical aperture (0.7) and then stitched them together. Second, the lung tissues exhibit high levels of autofluorescence which masks relatively weak smFISH signals. Autofluorescence is strongest in the green channels and diminishes at longer wavelengths. Therefore, we avoided the green and orange channels for smFISH. We reserved the green channel for antibody imaging, which yield signals strong enough to rise above the autofluorescence and the orange channel to acquire autofluorescence patterns (without using any probes or antibodies).

### Multiplexed analysis of rabbit granuloma with smFISH and immunofluorescence

We sought to perform smFISH in combination with traditional immunofluorescence (IF) as the latter can be used as a benchmark and to provide valuable controls. Given the pathogenicity of *Mtb*, it is necessary to formaldehyde fix the infected lung specimen under rather harsh conditions (10% formaldehyde for at least seven days). These conditions lead to sequestration of protein epitopes as well as RNAs and it is necessary to treat the FFPE sections with harsh steps, including boiling for antigen retrieval, which can lead to loss of RNA. We optimized this process using stringent RNase-free conditions that allow for antigen retrieval while preserving the RNA. We then performed IF against ionized calcium-binding adapter (IBA1), a protein marker for tissue-resident macrophages (19, 20), fixed the section again to ensure that the antibody will remain bound, and then performed smFISH using a set of Cy5-labeled probes (48 oligonucleotides) that were complementary to mRNA encoding rabbit IFNγ, which is a pro-inflammatory marker. The resulting images demonstrate the simultaneous detection of cells expressing a protein and a mRNA marker (Figure 1B). Although the well-segregated single molecule spots, characteristic of smFISH, were not visible in low magnification images, many such spots were visible in high resolution images, particularly in cells expressing IFNγ mRNA sparsely (Figure 1B). The images are consistent with expectation that some cells (macrophages) in the granuloma will express IBA1 alone, some will express IFNγ alone (T cells), and some will express both (activated macrophages), as alveolar macrophages are known to produce IFNγ avidly upon stimulation (21). The specificity of the anti-IBA1 antibody and IFNγ probes have been demonstrated earlier (4, 20).

### Identification of cell types based on combinatorial expressions of multiple markers

To demonstrate the detection of cell types by combinatorial expression of two mRNA and one protein marker, we included probes against TNFα mRNA (another pro-inflammatory marker) that were labeled with Texas Red in the experiment discussed above. A primary image showing the expression of the IBA1 protein and IFNγ and TNFα mRNAs is presented in Figure 2A.

To quantify the expression of each of the three markers in single cells within the section, we first computationally defined the locations and boundaries of cells in the section using DAPI staining of the nuclei (not shown in Figure 2A). The location and boundaries of DAPI stained nuclei were determined using a machine learning algorithm, StarDist, that can identify and demarcate the boundaries of nuclei even in a crowded spaces (22). The coordinates of the segmented nuclei were used to determine the approximate boundaries of cells around each nucleus by expanding the nuclear boundaries until they met with the cell boundaries of neighboring nuclei using QuPath (23). The accuracy of this method is depicted in Figure 2B (upper left panel). We then measured the cell intensity (mean pixel intensity within cell boundaries) in each of the three channels corresponding to IF for IBA1 and smFISH signals of TNFα and IFNγ mRNAs. Thereafter, we classified cells that expressed these markers above certain thresholds, separately for each of the three channels (Figure 2B). The thresholds were chosen interactively using cell intensity histograms such that most cells expressing the marker would be included and most cells not expressing would be excluded from the classifier. In addition to identifying cells that expressed the three markers individually, we also identified those that expressed them in various combinations. A digital map of the section was created by presenting each cell category using a distinct color. This map was overlayed on the primary image (Figure 2C). The number of cells in each category that we found in the section is presented in Figure 2D along with the color keys. By adjusting the opacity of the digital map, we could confirm that this map faithfully represents the underlying primary image for each of the three fluorescence channels.

Expression of various marker combinations indicates that the cells expressing just IBA1 are naïve macrophages, cell expressing IBA1 in combination with either or both cytokines are activated macrophages, cells not expressing IBA1 but expressing either or both cytokines are activated T cells, and cells expressing none of the markers are lung epithelial cells.

To define the microenvironments within granuloma where different cell types are concentrated, we created plots of density of the different cell types and overlaid them on the digital cell map (Figure 2E). These plots indicate that the solid core of this granuloma (which is a relatively young granuloma at four-week post infection) is largely populated by activated macrophages and activated T cells, whereas its periphery contains activated T cells, naïve macrophages, and lung epithelial cells. Multifunctional cells that express all three markers are restricted to the core of granuloma. Similar microenvironmental features have recently been observed MIBI-TOF imaging of human granulomas (13).

### mRNAs as surrogates for proteins

mRNAs have extensively been used as a surrogate for proteins in many RT-PCR, microarray and RNAseq studies. However, post-translational protein stabilization or decay (24) and stochastic mRNA synthesis (25) can create a discordance between mRNA and protein amounts in single cells. A particularly interesting case is hypoxia inducible factor-1α (HIF-1α) which is an oxygen sensing transcription factor that is degraded under normal oxygen concentrations but is stable under hypoxic conditions, such as those occurring during microbial infections (26). In addition to stabilizing the HIF-1α protein, microbial infections including *Mtb* infection (20), stimulate HIF-1α mRNA synthesis by producing transcription factor NF-κB which in turn induces transcription from the HIF-1α gene (27–30). We explored relative abundance of HIF-1α mRNA and protein within granuloma by imaging both simultaneously.

By imaging HIF-1α protein with an antibody and the HIF-1α mRNA with smFISH probes we found that about 10% of cells in rabbit granuloma expressed HIF-1α (Figure 3). As expected in the high magnification imaging (63x objective), smFISH signals were visible as discrete spots while the protein signals were more diffused. To explore the relative abundance of HIF-1α mRNA and proteins in these cells we segmented the cells using the combined sum of the fluorescence in the protein and mRNA channels rather than the DAPI fluorescence as was done in figure 2 and then determined the average pixel fluorescence within cell boundaries in each of the two channels. We found that out of 148 cells that we analyzed from three fields within one granuloma, 31 expressed only HIF-1α protein, 5 expressed only HIF-1α mRNA and 111 expressed both (Figure 3B). The mRNA and protein fluorescence signal in the cells that expressed both were correlated (correlation coefficient 0.66). Since in *Mtb* infected cells HIF-1α mRNA synthesis is stimulated and HIF-1α protein is stabilized (20, 27, 28), the correlation of the HIF-1α mRNA and protein in these cells is indicative of *Mtb* infection and also supports the idea that mRNA can serve as a reasonable surrogate for proteins.

### Detection of cells expressing transcripts at low levels using amplified single molecule FISH (ampFISH)

In the forgoing experiments cytokine mRNAs could be detected readily using smFISH because they are abundantly expressed in activated immune cells. However, we found that mRNAs encoding surface receptors, the traditional phenotypic markers of lymphocytes, yielded rather faint signals when low magnification was used. Therefore, we used ampFISH to demonstrate detection of mRNA encoding CD3ε (ε chain of cluster of differentiation 3) which serves as marker of T-cells since it is a component of the T cell receptor complex. In ampFISH a pair of hairpin probes is used whose binding at adjacent location on a target mRNA drives a conformational reorganization in one of the probes (Figure 4A) (18). This reorganization reveals a sequestered sequence which initiates a hybridization chain reaction (HCR), depositing multiple copies of fluorescent labels at the site. This signal amplification can be further increased by tiling multiple pairs of ampFISH probe across the length of a target mRNA (Figure 4B).

We designed 22 pairs of ampFISH probes complementary to mRNA encoding CD3ε and used them in combination with a set of 48 traditional smFISH probes against IFNγ mRNA. For these experiments we imaged an archived necrotic granuloma obtained from a Rhesus Macaque infected with *Mtb* (12). As mentioned before, these granulomas mimic human disease very closely. After hybridization with all the probes in a common hybridization mixture and removal of the excess probes by washing, HCR was performed to create ampFISH signal in a channel distinct from smFISH probe labels. Color combined images for the IFNγ smFISH and CD3ε ampFISH signals are presented in Figure 4C.

These images show single cells that express the two markers scattered around the periphery of a central necrotic region (Figure 4C). Inside the necrotic region, cells are present at various degrees of cellular degradation as indicated by a compression of nuclei in DAPI images (not shown) and loss of cellular morphology. Therefore, we did not use DAPI signals for the identification of cell boundaries as we did in the earlier experiment, instead we used the sum of smFISH and ampFISH signals as the basis of cell segmentation. After identifying all the cells that expressed both markers, we determined the fluorescence of each cell in both channels and classified them as either IFNγ+, CD3ε+, or both IFNγ+ and CD3ε+ using QuPath. The defined cells are indicated by colored circles (which are larger in size than the cells) that are overlayed on the primary image in Figure 4C and the distribution of their fluorescence is depicted in Figure 4D. Enlargements of a region of the granuloma, indicated by a blue square in Figure 4C, are also presented to demonstrate the accuracy of cell classification. In these enlargements, images from each of the two channels are presented in separate grey scale images with overlays indicating cell classifications. This colocalization analysis reveals that a small fraction of T cells (CD3ε+) also express IFNγ, indicating that they are in a state of activation. We found that only 6% of CD3ε+ cells were expressing IFNγ, which agrees with the flow cytometry data from dissociated cells obtained from macaque granulomas (31). The cells that express IFNγ but not CD3ε are likely to be predominantly activated macrophages.

### Iterative smFISH

Using probe sets labeled with distinguishable fluorophores, it is possible to image four mRNAs simultaneously in cultured cells. However, the autofluorescence in lung tissues limits us it further to only two or three clearly distinguishable fluorophores. A powerful strategy to achieve higher levels of multiplexing is to perform iterative hybridization in which multiple mRNAs are interrogated in successive cycles of binding target specific probes, imaging, and signal removal. Using this approach 33 mRNA species have successfully been imaged in brain sections (32). Two groups have achieved dramatically higher levels of multiplexing (100s to 1000s of target mRNA) by using pools of probes for all target mRNAs in a common hybridization reaction (MERFISH, (33) and seqFISH, (34)). ‘Primary target-binding’ probes are themselves not labeled with dyes but contain address tag sequences that in turn can be detected using secondary ‘readout probes’ (Figure 5). All target mRNAs can be detected by performing multiple cycles of imaging, where in each cycle, several readout probes labeled with distinguishable fluorophores are bound, followed by imaging and then dissociation of probes (Figure 5). High depths of multiplexing are achieved by combining multiple rounds of smFISH with color coding of probes, where each target is identified by a combination of colors rather than a unique color (33, 34). Given the limitations inherent in the *Mtb* granuloma imaging (high autofluorescence, heavy crosslinking, age of archived samples and low magnifications), we utilized a simplified version of iterative smFISH with a modest level of multiplexing, where pooled primary probes were used and multiple rounds of smFISH were performed, but no color coding was employed.

**Fig. 5.**
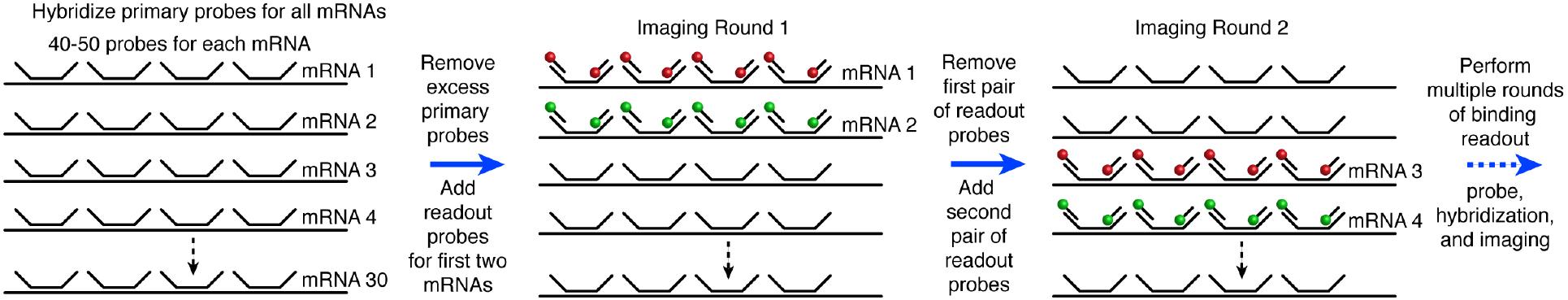
A schematic description of iterative hybridization for imaging many mRNA markers. A pool of primary probes against all mRNA target species is hybridized to the specimen overnight. After removal of excess primary probes, the mRNAs are identified, two at a time, by hybridizing a pair of ‘readout’ probes that bind to the unique address tags appended to the ends of each probe. All 48 probes for a given mRNA species have a unique address code. A series of four readout probe hybridization, imaging, and removal steps are performed to image eight mRNA species.

We prepared a pool of probes against eight different rabbit mRNAs that were likely to be expressed in that are likely to be expressed by myeloid cells in rabbit granulomas: NOS2, CD163, CD3, IL10, GZMH, S1008/9, IFNγ, and ARG1. The probe pool consisted of a total of 359 probes with 36-47 probes being complementary to one mRNA species (Supplementary Table 1). All probes for each mRNA species contained an address tag sequence appended to each end. The tag was unique for that mRNA species, but it was distinct from the address tags of other mRNA species. The address tags do not bind to the mRNA and remain single stranded when the target-specific region of the probe is bound to the mRNA. The presence of the address tag at the target could be determined by using fluorescently labeled ‘readout probes’ complementary to the address tags (Figure 5).

To ensure that the primary probes remain bound while repeated cycles of readout probe binding and removal are carried out, we designed relatively long (22-25 nucleotide (nt)) target specific regions in the primary probes and used 20 nt, long low melting temperature readout probes. The primary probes form RNA-DNA hybrids and the later a DNA-DNA which are inherently weaker. We included 10% formamide in the medium used to form the primary probe hybrids and 30% formamide in the medium to remove the readout probes. The 30% formamide destabilizes the weak hybrid between the readout probe and the address tag, but not between the primary probe and the target mRNA. To ensure that the readout probes would be able to bind rapidly (within 20-30 min), we used only three nucleotides in the design of the probes and used ethylene carbonate buffer for the readout probe binding. Each of these measures independently accelerates hybridization reactions (35–37). The readout probes were designed to detect only its cognate address tag and not cross react with the other address tags. This feature of the readout probe set was confirmed by *in vitro* fluorescence DNA melting assays performed before their deployment *in situ*.

The pool of mRNA specific primary probes was hybridized to the tissue sections in an overnight hybridization reaction. After removing the excess primary probes, we interrogated the eight target mRNAs in four successive rounds of hybridization and imaging with pairs of labeled readout probes. In each round, one Texas Red labeled readout probe complementary to the address tag of the first mRNA and another Cy5 labeled readout probe which was specific to the address tag of the second mRNA were hybridized to the sample. After imaging the first pair of mRNA targets in each channel, the first pair of readout probes was removed and the readout probes for the second set of mRNA targets were added. Four rounds of hybridization with a total of eight readout probes yielded images for all eight mRNA targets (Figure 6).

**Fig. 6.**
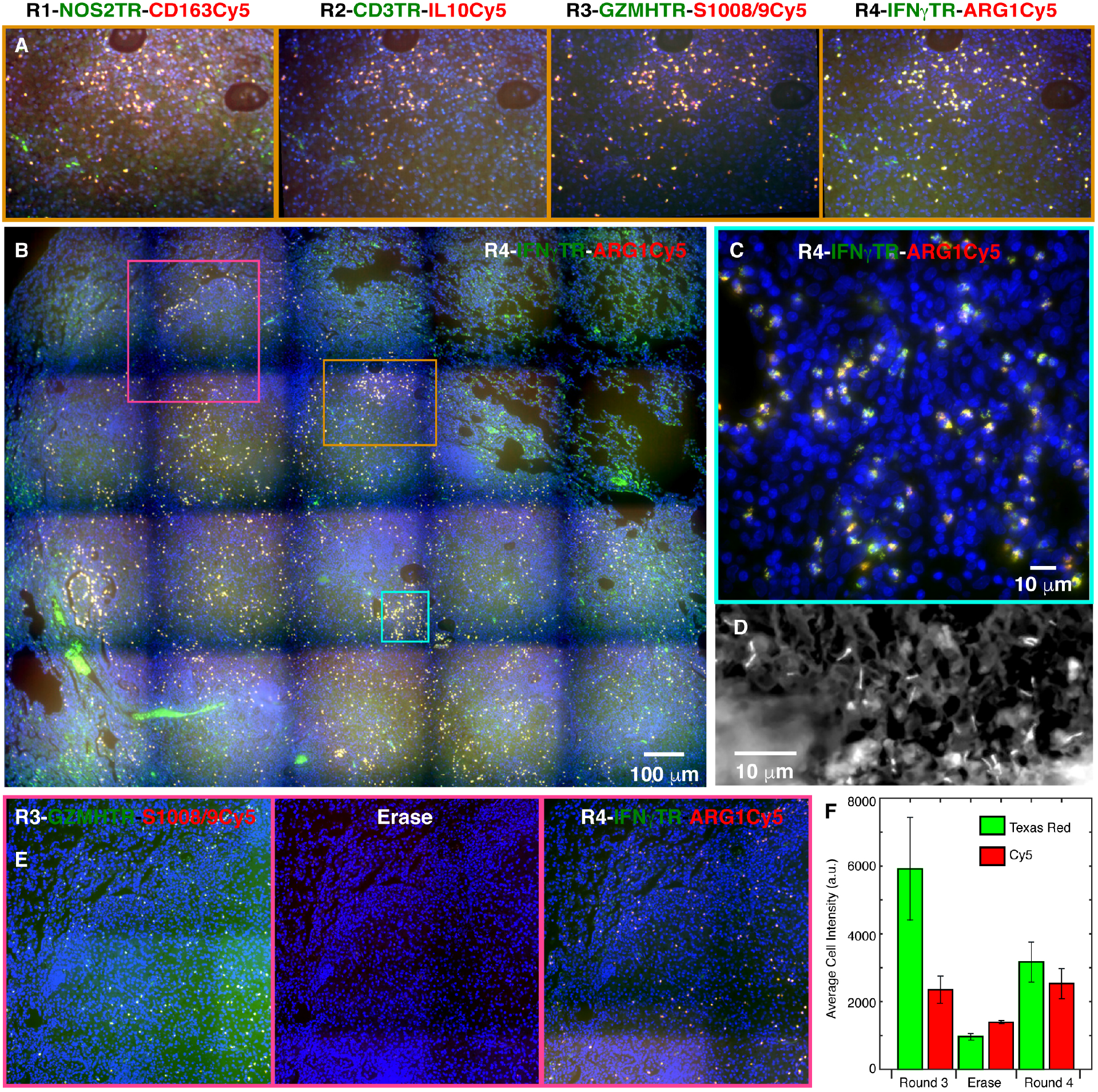
Demonstration of imaging cells expressing eight different mRNA targets in a solid rabbit granuloma. The color coding in each image is identified by the color of RNA labels (blue is DAPI). Montages like the one shown in panel **B** (which is for round 4) were obtained using a 20x objective in each of the four rounds of iterative smFISH and then stitched together. Panels in **A** show the images obtained in each round for the region demarcated by the orange square. Panel **C** represents a high magnification (63x objective) image from cyan square. Panel **D** shows detection of *Mtb* in the granuloma with rhodamine-auramine staining performed after round four. Panels in **E**, which correspond to purple square, demonstrate that the signal in round 3 is lost upon removal of readout probes and new signals appear after addition of round 4 readout probes. Graph **F** shows the fluorescence intensities in single cells for two markers in round 3, erase step, and round 4. After an erase step we are able to successfully re-image mRNAs seen in any of the previous round without a significant loss of signal.

Each round of imaging lit up a specific set of cells in the granuloma with most cells exhibiting little or no background. We found that many cells expressed multiple markers although the relative expression levels of different markers varied from cell to cell. High magnification imaging of a portion of the granuloma in round four, which imaged IFNγ and ARG1 mRNAs, exhibited spot-like signals that are characteristic of smFISH. This image shows that while many cells express both markers, some exhibit only IFNγ signal (Figure 6C) and points to the specificity of each probe set in the pair and to the diversity of cell types in this milieu.

To show that the readout probes are removed while the primary probes remain bound in each round, we imaged the section after the third round without adding a readout probe set. This experiment showed that the cellular fluorescence intensity is reduced to background levels after removal of the readout probes and becomes high again in the fourth round (Figures 6E&F).

Finally, exploiting the open-dish imaging format, we performed rhodamine-auramine staining for the detection of *Mtb* from the same section after completion of the iterative FISH (Figure 6D).

## Discussion

Here we demonstrate imaging of many mRNA species expressed in immune cells in archived tuberculosis granulomas obtained from rabbits and macaques. These results indicate that despite the challenges of autofluorescence, extensive formaldehyde crosslinking and the age of the specimens, sufficient mRNAs survive to enable their detection by smFISH. In the case of sparsely expressed mRNAs we utilized ampFISH which could be performed in multiplex with the conventional smFISH.

Although the general specificity of smFISH and ampFISH has been establihed earlier (15, 18, 38), our results show that the targeted mRNAs were accurately detected. Particularly, the same cells showed the presence of HIF-1α mRNA and protein (Figure 3), and the percent of CD3ε+ cells that expressed IFNγ matched with earlier flow cytometric studies of T cells in cells dissociated from macaque granuloma (Figure 4).

An important aspect of this study is the digital identification of cells based on combination of markers that they express. We segmented cells using a machine learning algorithm, StarDist implemented in the QuPath environment. This digital identification of cells based on cell markers allows unbiased quantitative morphometric analysis, such as microenvironment analysis through cell type density maps, which will allow unbiased explorations of cell-to-cell interactions which give rise to structures like granulomas.

Granulomas harbor a great variety of immune cells including macrophages differentiated along a spectrum of pro- and anti-inflammatory states, and T cells present in various stages of activation and differentiation (2, 39). Furthermore, the observations that macrophages, dendritic cells and T cells undergo shifts in their metabolic program upon encountering pathogens (40–42) point to additional diversity of cell states. Classical phenotypic surface markers are not sufficient to capture this diversity. However, with our platform it will be possible to obtain sets of suitable mRNA markers for cell subtype identification from single cell RNA seq data and then use them to image the desired cell subtypes *in situ*. Furthermore, it will be possible to perform such analyses on human granulomas when they are accessible.

We obtained a depth of multiplexing of eight mRNAs by performing four rounds of iterative hybridization using readout probes in two colors on archived tissues from *Mtb* infected animals. This can easily be extended to about 30 species of mRNA by performing 15 rounds of hybridization. This throughput is still modest compared to 1000s of mRNA species that recent spatial transcriptomic technologies are able to achieve (43). To achieve that level of throughput, it will be necessary to use four colors per round of hybridization, implement color coding schemes for target detection, and embed sections into acrylamide gels followed by clearing. However, to perform all these steps in a biosafety level 3 environment on fresh *Mtb* infected tissue is currently challenging.

In conclusion, we describe powerful methods for future studies of granuloma architecture to understand how sometimes the tussle between the pro- and anti-inflammatory activities of various cell types within granulomas leads to sequestration of *Mtb* into long-term non-replicating latency, whereas, at other times it leads to dissemination of replicating bacteria into the larger tissue, causing active disease.

## Supporting information

Supplementary tabl1 1

## Acknowledgements

The work was supported by National Institutes of Health (NIH) grants R01AI127844 to L.S. and S.S., R21AI163824 to L.S. and R01 CA227291 to S.T.. Additionally, a Bill and Melinda Gates Foundation grant OPP1157210, provided support to S.S and a New Jersey Health Foundation Grant to S.T. We thank Ryan Dikdan for his careful reading of the manuscript.

## Notes

### Competing Interest Statement

Rutgers University receives royalties from the sale of prelabeled sm-FISH probes by LGC Biosearch Technologies, which markets them as Stellaris probes. A portion of these proceeds is distributed to S.T.'s laboratory for research support and to him personally. These proceeds do not influence the conclusions of this research.

## Bibliography

1. Ramakrishnan, L. 2012. Revisiting the role of the granuloma in tuberculosis. Nat Rev Immunol 12: 352–366.

2. Cadena, A. M., S. M. Fortune, and J. L. Flynn. 2017. Heterogeneity in tuberculosis. Nat Rev Immunol 17: 691–702.

3. Flynn, J. L., and J. Chan. 2022. Immune cell interactions in tuberculosis. Cell 185: 4682–4702.

4. Bushkin, Y., F. Radford, R. Pine, A. Lardizabal, B. T. Mangura, M. L. Gennaro, and S. Tyagi. 2015. Profiling T cell activation using single-molecule fluorescence in situ hybridization and flow cytometry. J Immunol 194: 836–841.

5. IGRA_Assays. 2020. https://www.cdc.gov/tb/publications/factsheets/testing/igra.htm.

6. Berry, M. P., C. M. Graham, F. W. McNab, Z. Xu, S. A. Bloch, T. Oni, K. A. Wilkinson, R. Banchereau, J. Skinner, R. J. Wilkinson, C. Quinn, D. Blankenship, R. Dhawan, J. J. Cush, A. Mejias, O. Ramilo, O. M. Kon, V. Pascual, J. Banchereau, D. Chaussabel, and A. O’Garra. 2010. An interferon-inducible neutrophil-driven blood transcriptional signature in human tuberculosis. Nature 466: 973–977.

7. Kaplan G, T., L. Tsenova. 2011. Pulmonary tuberculosis in the rabbit. A color atlas of comparative pathology of pulmonary tuberculosis. only F. J. Leong, Dartois V, Dick T, ed, Boca Raton, FL. 107–130.

8. Driver, E. R., G. J. Ryan, D. R. Hoff, S. M. Irwin, R. J. Basaraba, I. Kramnik, and A. J. Lenaerts. 2012. Evaluation of a mouse model of necrotic granuloma formation using C3HeB/FeJ mice for testing of drugs against Mycobacterium tuberculosis. Antimicrob Agents Chemother 56: 3181–3195.

9. Carow, B., T. Hauling, X. Qian, I. Kramnik, M. Nilsson, and M. E. Rottenberg. 2019. Spatial and temporal localization of immune transcripts defines hallmarks and diversity in the tuberculosis granuloma. Nat Commun 10: 1823.

10. Subbian, S., L. Tsenova, M. J. Kim, H. C. Wainwright, A. Visser, N. Bandyopadhyay, J. S. Bader, P. C. Karakousis, G. B. Murrmann, L. G. Bekker, D. G. Russell, and G. Kaplan. 2015. Lesion-Specific Immune Response in Granulomas of Patients with Pulmonary Tuberculosis: A Pilot Study. PLoS One 10: e0132249.

11. Mattila, J. T., O. O. Ojo, D. Kepka-Lenhart, S. Marino, J. H. Kim, S. Y. Eum, L. E. Via, C. E. Barry 3rd, E. Klein, D. E. Kirschner, S. M. Morris, Jr., P. L. Lin, and J. L. Flynn. 2013. Microenvironments in tuberculous granulomas are delineated by distinct populations of macrophage subsets and expression of nitric oxide synthase and arginase isoforms. J Immunol 191: 773–784.

12. Mehra, S., B. Pahar, N. K. Dutta, C. N. Conerly, K. Philippi-Falkenstein, X. Alvarez, and D. Kaushal. 2010. Transcriptional reprogramming in nonhuman primate (rhesus macaque) tuberculosis granulomas. PLoS One 5: e12266.

13. McCaffrey, E. F., M. Donato, L. Keren, Z. Chen, A. Delmastro, M. B. Fitzpatrick, S. Gupta, N. F. Greenwald, A. Baranski, W. Graf, R. Kumar, M. Bosse, C. C. Fullaway, P. K. Ramdial, E. Forgo, V. Jojic, D. Van Valen, S. Mehra, S. A. Khader, S. C. Bendall, M. van de Rijn, D. Kalman, D. Kaushal, R. L. Hunter, N. Banaei, A. J. C. Steyn, P. Khatri, and M. Angelo. 2022. The immunoregulatory landscape of human tuberculosis granulomas. Nat Immunol 23: 318–329.

14. Femino, A. M., F. S. Fay, K. Fogarty, and R. H. Singer. 1998. Visualization of single RNA transcripts in situ. Science 280: 585–590.

15. Raj, A., P. van den Bogaard, S. A. Rifkin, A. van Oudenaarden, and S. Tyagi. 2008. Imaging individual mRNA molecules using multiple singly labeled probes. Nat Methods 5: 877–879.

16. Subbian, S., L. Tsenova, G. Yang, P. O’Brien, S. Parsons, B. Peixoto, L. Taylor, D. Fallows, and G. Kaplan. 2011. Chronic pulmonary cavitary tuberculosis in rabbits: a failed host immune response. Open Biol 1: 110016.

17. Raj, A., and S. Tyagi. 2010. Detection of individual endogenous RNA transcripts in situ using multiple singly labeled probes. Methods Enzymol 472: 365–386.

18. Marras, S. A. E., Y. Bushkin, and S. Tyagi. 2019. High-fidelity amplified FISH for the detection and allelic discrimination of single mRNA molecules. Proc Natl Acad Sci U S A 116: 13921–13926.

19. Imai, Y., I. Ibata, D. Ito, K. Ohsawa, and S. Kohsaka. 1996. A novel gene iba1 in the major histocompatibility complex class III region encoding an EF hand protein expressed in a monocytic lineage. Biochem Biophys Res Commun 224: 855–862.

20. Shi, L., H. Salamon, E. A. Eugenin, R. Pine, A. Cooper, and M. L. Gennaro. 2015. Infection with Mycobacterium tuberculosis induces the Warburg effect in mouse lungs. Sci Rep 5: 18176.

21. Darwich, L., G. Coma, R. Pena, R. Bellido, E. J. Blanco, J. A. Este, F. E. Borras, B. Clotet, L. Ruiz, A. Rosell, F. Andreo, R. M. Parkhouse, and M. Bofill. 2009. Secretion of interferon-gamma by human macrophages demonstrated at the single-cell level after costimulation with interleukin (IL)-12 plus IL–18. Immunology 126: 386–393.

22. Schmidt, U., Weigert, M, Broaddus, C, Myers, G. 2018. Cell Detection with Star-convex Polygons. In International Conference on Medical Image Computing and Computer-Assisted Intervention (MICCAI), Granada, Spain, September 2018.

23. Bankhead, P., M. B. Loughrey, J. A. Fernandez, Y. Dombrowski, D. G. McArt, P. D. Dunne, S. McQuaid, R. T. Gray, L. J. Murray, H. G. Coleman, J. A. James, M. Salto-Tellez, and P. W. Hamilton. 2017. QuPath: Open source software for digital pathology image analysis. Sci Rep 7: 16878.

24. Liu, Y., A. Beyer, and R. Aebersold. 2016. On the Dependency of Cellular Protein Levels on mRNA Abundance. Cell 165: 535–550.

25. Raj, A., C. S. Peskin, D. Tranchina, D. Y. Vargas, and S. Tyagi. 2006. Stochastic mRNA synthesis in mammalian cells. PLoS Biol 4: e309.

26. Ivan, M., K. Kondo, H. Yang, W. Kim, J. Valiando, M. Ohh, A. Salic, J. M. Asara, W. S. Lane, and W. G. Kaelin, Jr. 2001. HIFalpha targeted for VHL-mediated destruction by proline hydroxylation: implications for O2 sensing. Science 292: 464–468.

27. Rius, J., M. Guma, C. Schachtrup, K. Akassoglou, A. S. Zinkernagel, V. Nizet, R. S. Johnson, G. G. Haddad, and M. Karin. 2008. NF-kappaB links innate immunity to the hypoxic response through transcriptional regulation of HIF–1alpha. Nature 453: 807–811.

28. Osada-Oka, M., N. Goda, H. Saiga, M. Yamamoto, K. Takeda, Y. Ozeki, T. Yamaguchi, T. Soga, Y. Tateishi, K. Miura, D. Okuzaki, K. Kobayashi, and S. Matsumoto. 2019. Metabolic adaptation to glycolysis is a basic defense mechanism of macrophages for Mycobacterium tuberculosis infection. Int Immunol 31: 781–793.

29. Nizet, V., and R. S. Johnson. 2009. Interdependence of hypoxic and innate immune responses. Nat Rev Immunol 9: 609–617.

30. Shi, L., E. A. Eugenin, and S. Subbian. 2016. Immunometabolism in Tuberculosis. Front Immunol 7: 150.

31. Gideon, H. P., J. Phuah, A. J. Myers, B. D. Bryson, M. A. Rodgers, M. T. Coleman, P. Maiello, T. Rutledge, S. Marino, S. M. Fortune, D. E. Kirschner, P. L. Lin, and J. L. Flynn. 2015. Variability in tuberculosis granuloma T cell responses exists, but a balance of pro- and anti-inflammatory cytokines is associated with sterilization. PLoS Pathog 11: e1004603.

32. Codeluppi, S., L. E. Borm, A. Zeisel, G. La Manno, J. A. van Lunteren, C. I. Svensson, and S. Linnarsson. 2018. Spatial organization of the somatosensory cortex revealed by osmFISH. Nat Methods 15: 932–935.

33. Chen, K. H., A. N. Boettiger, J. R. Moffitt, S. Wang, and X. Zhuang. 2015. RNA imaging. Spatially resolved, highly multiplexed RNA profiling in single cells. Science 348: aaa6090.

34. Eng, C. L., M. Lawson, Q. Zhu, R. Dries, N. Koulena, Y. Takei, J. Yun, C. Cronin, C. Karp, G. C. Yuan, and L. Cai. 2019. Transcriptome-scale super-resolved imaging in tissues by RNA seqFISH. Nature 568: 235–239.

35. Zhang, Z., A. Revyakin, J. B. Grimm, L. D. Lavis, and R. Tjian. 2014. Single-molecule tracking of the transcription cycle by sub-second RNA detection. Elife 3: e01775.

36. Matthiesen, S. H., and C. M. Hansen. 2012. Fast and non-toxic in situ hybridization without blocking of repetitive sequences. PLoS One 7: e40675.

37. Moffitt, J. R., J. Hao, G. Wang, K. H. Chen, H. P. Babcock, and X. Zhuang. 2016. High-throughput single-cell gene-expression profiling with multiplexed error-robust fluorescence in situ hybridization. Proc Natl Acad Sci U S A 113: 11046–11051.

38. Batish, M., P. van den Bogaard, F. R. Kramer, and S. Tyagi. 2012. Neuronal mRNAs travel singly into dendrites. Proc Natl Acad Sci U S A 109: 4645–4650.

39. Mosser, D. M. 2003. The many faces of macrophage activation. J Leukoc Biol 73: 209–212.

40. Rodriguez-Prados, J. C., P. G. Traves, J. Cuenca, D. Rico, J. Aragones, P. Martin-Sanz, M. Cascante, and L. Bosca. 2010. Substrate fate in activated macrophages: a comparison between innate, classic, and alternative activation. J Immunol 185: 605–614.

41. Krawczyk, C. M., T. Holowka, J. Sun, J. Blagih, E. Amiel, R. J. DeBerardinis, J. R. Cross, E. Jung, C. B. Thompson, R. G. Jones, and E. J. Pearce. 2010. Toll-like receptor-induced changes in glycolytic metabolism regulate dendritic cell activation. Blood 115: 4742–4749.

42. Xu, K., N. Yin, M. Peng, E. G. Stamatiades, A. Shyu, P. Li, X. Zhang, M. H. Do, Z. Wang, K. J. Capistrano, C. Chou, A. G. Levine, A. Y. Rudensky, and M. O. Li. 2021. Glycolysis fuels phosphoinositide 3-kinase signaling to bolster T cell immunity. Science 371: 405–410.

43. Tian, L., F. Chen, and E. Z. Macosko. 2022. The expanding vistas of spatial transcriptomics. Nat Biotechnol.

